# CCR2 identifies tendon resident macrophage and T cell populations while CCR2 deficiency impedes late tendon healing

**DOI:** 10.1101/2022.07.20.500814

**Authors:** Samantha Muscat, Anne E.C. Nichols, Emma Gira, Alayna E. Loiselle

**Author notes:** <u>Corresponding Author</u> Alayna E. Loiselle, PhD, Center for Musculoskeletal Research, University of Rochester Medical Center, 601 Elmwood Ave, Box 665-G, Rochester, NY, 14642.

## Abstract

During tendon healing, macrophages are thought to be a key mediator of scar tissue formation, which prevents successful functional restoration of the tendon. However, macrophages are critical for successful tendon healing as they aid in wound debridement, extracellular matrix deposition, and promote fibroblast proliferation. Recent work has sought to better define the multi-faceted functions of macrophages using depletion studies, while other studies have identified a tendon resident macrophage population. To begin to delineate the functions of tendon-resident versus circulation-derived macrophages, we examined the tendon healing phenotype in Chemokine Receptor 2 (CCR2) reporter (CCR2^GFP/+^), and knockout mice. CCR2 is a chemokine receptor primarily found on the surface of circulating bone marrow derived monocytes, with CCR2 being an important mediator of macrophage recruitment to wound environments. Surprisingly, CCR2^GFP/+^ cells were present in the tendon during adult homeostasis, and single cell RNA sequencing identified these cells as tendon-resident macrophages and T cells. During both homeostasis and healing, CCR2 knockout resulted in a substantial decrease in CCR2^GFP+^ cells and pan-macrophages. Additionally, loss of CCR2 resulted in reduced numbers of myofibroblasts and impeded functional recovery during late healing. This study highlights the heterogeneity of tendon-resident and recruited immune cells and their contributions following injury, and establishes an important role for CCR2 in modulating both the adult tendon cell environment and tendon healing process.

## INTRODUCTION

Tendons are dense connective tissues that are primarily composed of type I collagen. Once injured, tendon function is never fully restored. Instead, tendons heal with the formation of a fibrovascular scar, characterized by excessive and disorganized extracellular matrix (ECM) deposition(1,2). While this scar tissue provides tissue continuity and a degree of stability, the mechanical composition of the tendon is compromised, increasing risk of re-rupture(3). Recent work has suggested that macrophages are a key driver of this scar tissue response both in tendon(1,2,4,5) and other tissues(6,7).

While tissue resident macrophages are thought to be a relatively small population of the tendon cell environment during homeostasis(8), there is a robust increase in extrinsic macrophage infiltration to the tendon after acute injury, which is critical for initiation of the inflammatory phase of healing(1,4,5,9). Macrophages are a dynamic cell population that interact with tissue resident cell populations, extrinsic immune cells, and extracellular proteins(6,7,10). Following tissue injury, apoptotic cells release damage-associated molecular pattern (DAMPS) that activate pathways in macrophages that recruit neutrophils, monocytes, and other inflammatory cells to the injured tissue. Macrophages also secrete anti-inflammatory molecules, growth factors, and matrix metalloproteinases that promote tissue repair and matrix degradation. Macrophages are also known to create a pro-fibrotic feedback loop with myofibroblasts, a specialized contractile fibroblast that drives both physiological healing and the transition to fibrosis(11-13). The diverse function of macrophages allows them to successfully evade pathogens, clear wound debris and deposit collagen in an ever-changing microenvironment. However, failure to clear macrophages following the acute inflammatory phase is associated with a transition to a more chronic, fibrotic phenotype resulting in excessive scar tissue deposition.

In contrast to other tissues, the contributions of macrophages during tendon healing are not well defined. Given the fibrotic healing process that occurs in tendon and the intricate link between macrophages and fibrosis, ascertaining the role of macrophages during tendon healing is therefore critical to identifying potential therapeutic approaches. Recent work has utilized macrophage depletion approaches to better understand the role of macrophages during tendon healing. Macrophage depletion in adult tendon promoted cell proliferation and ECM accumulation but lead to inferior tensile strength(14). Consistent with this, macrophage depletion in neonatal Achilles tendons resulted in impaired functional healing, however cell proliferation was reduced(9). In addition, extracellular vesicles derived from human mesenchymal stem cells (MSCs) primed macrophages towards an M2 anti-inflammatory phenotype that improved Achilles tendon biomechanics following injury(15). However, there are still important questions regarding the potentially unique roles of tissue-resident vs. circulation derived macrophages during tendon healing. Therefore, we sought to better understand the contribution of extrinsic macrophages during flexor tendon healing, using Chemokine Receptor 2 (CCR2)-reporter and CCR2 knock-out (CCR2^KO^) mice. CCR2 is a chemokine receptor primarily found on the surface of bone marrow derived monocytes circulating in the peripheral blood, with CCR2 thought to mediate recruitment of macrophages to the wound environment (16-18). While the role of CCR2 during tendon healing has yet to be investigated, it has been studied extensively in a variety of different tissues. For example, CCR2^KO^ kidneys had impaired macrophage and fibroblast accumulation during renal fibrosis(19) and CCR2^KO^ liver had reduced macrophage content concomitant with delayed liver fibrosis(20). In the present study, we defined the functional effects of CCR2 inhibition and tracked changes in CCR2+ cell recruitment during tendon healing. We hypothesized that following acute tendon injury, CCR2^KO^ would reduce recruitment of extrinsic macrophages and that diminished macrophage recruitment would subsequently impair myofibroblast differentiation and functional recovery.

## METHODS

### Animal Ethics

This study was carried out in strict accordance with the recommendations in the *Guide for the Care and Use of Laboratory Animals* of the National Institutes of Health (Bethesda, MD, USA). All animal procedures were approved by the University Committee on Animal Research (UCAR) at the University of Rochester (UCAR Number: 2014-004E).

### CCR2^GFP^ mouse model

CCR2^GFP/+^ mice were obtained from Jackson Laboratories (strain #027619). These transgenic mice have the green fluorescent protein (GFP) sequence followed by a polyadenylation signal inserted into the chemokine receptor 2 (CCR2) gene(21). In CCR2^GFP/+^ mice, one copy of CCR2 has been replaced with GFP, however mice are phenotypically normal and are used as a CCR2 reporter. In CCR2^KO^ mice, GFP has replaced both copies of CCR2, resulting in CCR2 knockout and functional impairment in extrinsic monocyte/ macrophage recruitment(21). CCR2^WT^ mice have normal CCR2 expression and do not express GFP. CCR2^GFP/+^ mice were bred to obtain CCR2^GFP/+^ reporter mice, knockout mice (CCR2^GFP/GFP;^ CCR2^KO^) and CCR2^WT^ controls.

### Flexor tendon repair

At 10-12 weeks of age, CCR2^WT^, CCR2^GFP/+^ and CCR2^KO^ mice underwent complete transection and repair of the flexor digitorum longus (FDL) tendon in the hind-paw as previously described(22). Briefly mice were given a pre-surgical dose of sustained release buprenorphine (0.5-1.0 mg/kg). Mice were then anesthetized with ketamine (60 mg/kg) and xylazine (4 mg/kg). Following preparation and sterilization of the surgical site, the FDL was surgically transected in the transverse plane at the myotendinous junction in the calf to prevent early-stage strain-induced rupture of the repair site. The skin was closed with a 5-0 suture. A small incision was then made on the posterior surface of the hind-paw, the FDL tendon was located and completely transected using micro spring scissors. The tendon was then repaired using 8-0 suture and the skin was closed with a 5-0 suture. Following surgery, mice were allowed to resume normal cage activity with regular food intake and water consumption.

### Histology and immunofluorescence

Following sacrifice, the hind paws of CCR2^WT^, CCR2^GFP/+^ and CCR2^KO^ were harvested for frozen sectioning. Hind paws were harvested at 8-,14-, 21-, and 28-days post-repair (n=3-5 per genotype per timepoint). Injured and contralateral hind paws were then fixed in 10% neutral buffer formalin for 24 hours at 4° C, decalcified in Webb-Jee 14% EDTA solution for 5 days at 4° C, and processed in 30% sucrose for 24 hours at 4° C to cryo-protect the tissue. Samples were then embedded in Cryomatrix (Thermo Fisher Scientific, Waltham, MA, USA) and sectioned into 8µm sagittal sections using a cryotape-transfer method(23). Sections were mounted on glass slides using 1% chitosan in 0.25% acetic acid. Slides were fixed and probed with antibodies for F4/80 (1:100 Abcam, #ab6640), CX3CR1 (1:200, Abcam, #ab8021), αSMA-Cy3 (1:250, Sigma-Aldrich, #C6198), and fibroblast activation protein (1:500, Abcam, #ab53066) and counterstained with NucBlue Live Cell Stain (Invitrogen, #R37605). Endogenous fluorescence was imaged with a VS120 Virtual Slide Microscope (Olympus, Waltham, MA).

### Quantification of fluorescence

Fluorescent images were processed using Visiopharm image analysis software v.6.7.9.2590 (Visiopharm, Horsholm, Denmark). Regions of interest (ROI) were drawn to include the tendon stubs and bridging scar tissue for processing. For CCR2^GFP^ staining, automatic segmentation using a threshold classifier was used to define discrete cell populations based on fluorescent intensity and data are given as a percentage of all cells within an ROI. For F4/80 and αSMA staining, the area of each fluorescent signal was calculated and data are presented as percent positive staining normalized to total area. An n = 3-5 for CCR2^WT^, CCR2^GFP/+^, CCR2^KO^ mice was used for quantification.

### Assessment of Gliding Function and biomechanical testing

Tendon gliding function was assessed as previously described(24-26). Briefly, injured and uninjured contralateral hindlimbs were harvested at the knee joint, and skin was removed down to the ankle. The end of the tendon was released just proximal to the tarsal tunnel without disrupting the skin or the ankle. The tendon end was secured using cyanoacrylate between two pieces of tape. The tibia was then secured in an alligator clip and loaded incrementally with small weights ranging from 0-19g. Digital images were taken at each weight. The flexion angle of the metatarsophalangeal (MTP) joint was measured from these images. The MTP flexion angle corresponds to the degrees of flexion upon application of the 19g weight. Uninjured tendons undergo complete flexion of the digits at 19g. Gliding resistance is based on the changes in MTP flexion angle of the range of the applied weights. Higher gliding resistance is indicative of impaired gliding function and the formation of adhesive scar tissue between the tendon and surrounding tissue.

Following gliding testing, the tibia was removed at the ankle and the toes and proximal section of the FDL in the tape were secured in opposing ends of custom grips on an Instron 8841 DynaMight™ axial servo hydraulic testing system (Instron Corporation, Norwood, MA). The tendons were loaded at a rate of 30 mm/minute until failure to test tensile strength. Force-displacement curves were plotted to determine maximum tensile force and stiffness. An n = 9-12 per genotype per time point were analyzed for both injured and contralateral tendons.

### Statistical Analysis

Quantitative data were analyzed in GraphPad Prism and are presented as means ± standard deviation. Normality was assessed using the Shapiro-Wilk normality test for all data. A student’s t test was used to assess differences in the number of CCR2^GFP^ cells between genotypes for uninjured data. A two-way Analysis of Variance (ANOVA) with Sidak’s multiple comparisons test was used to determine significance between genotypes at any given timepoint for injured quantification data and functional analysis (gliding and biomechanics). GraphPad Prism was used to detect statistical outlier data points (ROUT method, Q value= 1%). One outlier was identified in the D28 CCR2^GFP^+ cell quantification data set and was removed from analysis. For all experiments an n=1 represents one mouse. For all outcome measures, p ≤ 0.05 was considered significant. Significance is noted in all figures with the following conventions: *p≤0.05, **p≤0.01, ***p≤0.001, ****p≤0.0001.

## RESULTS

### CCR2 identifies tendon-resident cell populations

Given that CCR2+ cells are typically considered to be recruited from systemic circulation, we first determined whether any CCR2+ cells were present during adult tendon homeostasis using CCR2^GFP/+^ reporter mice. Surprisingly, CCR2+ cells accounted for 7.77 ± 5.372% of the total resident tendon cell population in CCR2^GFP/+^ mice (Figure 1A & B).

**Fig 1.**
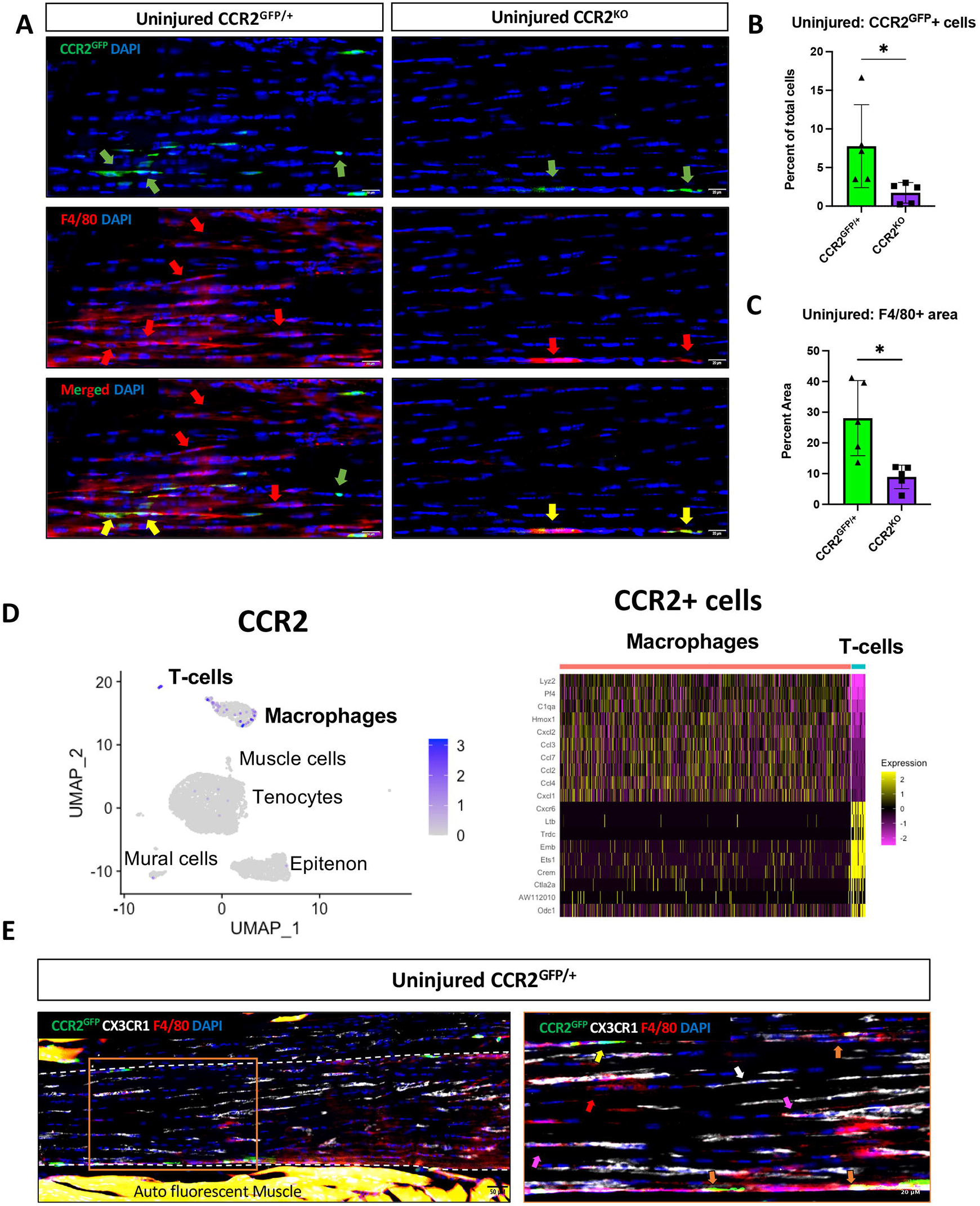
CCR2 identifies tendon-resident macrophages and T cells. (A) To assess the effects of CCR2 deficiency during tendon homeostasis, uninjured CCR2^GFP/+^ and CCR2^KO^ tendons were probed for GFP (CCR2^GFP)^) and F4/80. Relative to the CCR2^GFP/+^ control, CCR2^KO^ tendons have reduced presence of CCR2^GFP^+ cells (green arrows), F4/80+ macrophages (red arrows), double-positive cells (yellow arrows) and CCR2^GFP^+ F4/80- cells (green arrows). (B) Quantification of % CCR2^GFP^+ cells and (C) % area of F4/80+ macrophages in uninjured CCR2^GFP/+^ and CCR2^KO^ tendons. (D) UMAP from uninjured WT C57BL/6J tendons of CCR2+ cells, heatmap of CCR2+ expression in macrophage and T-cell clusters. (E) To examine the relationship between CCR2 and CXRCR1, CCR2^GFP/+^ reporter mice were stained for GFP (CCR2^GFP^), F4/80 and CX3CR1. Immunofluorescence exhibits 5 cell populations: CCR2^GFP^+ CX3CR1+ F4/80+ macrophages (orange arrows), CCR2^GFP^+ CX3CR1- F4/80+ (yellow arrows), CCR2^GFP^- CX3CR1+ F4/80+ macrophages (pink arrows) and CCR2^GFP^- CX3CR1- F4/80+ macrophages (red arrows) and CCR2^GFP^- CX3CR1+ F4/80- cells (white arrows). Nuclear live cell stain is blue. Tendon is outlined in white dotted line. Scale bars, 20µM and 50 µM. N=5 per genotype. Student’s *t* test was used to assess statistical significance. *p≤0.05.

However, in CCR2^KO^ tendons, only 1.73 ± 1.33% of the cells were GFP+ (Figure 1A & B). Collectively, these data suggest that CCR2 is required for incorporation of most CCR2^GFP^ cells into the native tendon, but that there are also CCR2-independent mechanisms responsible for the presence of CCR2+ cells during adult tendon homeostasis.

We next examined the relationship between tendon resident CCR2^GFP^ cells and the pan-macrophage marker F4/80. The majority of CCR2^GFP^+ cells were F4/80+ (Figure 1A & C; yellow arrows); however, there were also CCR2^GFP^- F4/80+ macrophages (Figure 1A; red arrows). With loss of CCR2, both of these populations are reduced, relative to CCR2^GFP/+^ controls (Figure 1A; yellow arrows). We also observed a significant 77.75% reduction in overall F4/80 surface-area staining in CCR2^KO^ tendons (Figure 1C; p = 0.0451) compared to the CCR2^GFP/+^ tendons. CCR2^KO^ tendons had an average of 8.92 ± 3.83% F4/80 area staining, compared to 28.07 ± 12.23% F4/80 area staining in the CCR2^GFP/+^ tendons (Figure 1A & C; red arrows). We also identified a CCR2^GFP^+ F4/80- resident cell population (Figure 1A; green arrow). Interestingly, these reductions in the uninjured CCR2^KO^ tendon are restricted to the mid-substance of the tendon, with the remaining CCR2^GFP^+ cells and F4/80 macrophages in the epitenon region. Taken together, these data indicate that CCR2 identifies a subset of tendon-resident macrophages in addition to a non-macrophage cell population in uninjured tendons, and that loss of CCR2 decreases the presence of both populations.

To further validate that CCR2 identifies a tendon-resident macrophage population and to better define the CCR2+ F4/80- population, we analyzed single cell RNA sequencing data from uninjured FDL tendons (full study to be published elsewhere). Consistent with our immunofluorescence staining, CCR2 expression was largely restricted to the tendon-resident macrophage population (Figure 1D). Interestingly, we also found that CCR2 was expressed by a tendon-resident T cell population (Figure 1D). While we initially hypothesized that CCR2 regulates extrinsic macrophage populations, these substantial alterations in the resident tendon cell environment in CCR2^KO^ tendons identifies a potential role for CCR2 in mediating the composition of the tendon intrinsic cell environment as well.

A tendon resident macrophage population has been previously defined as CX3CR1+(8). Thus, we stained uninjured CCR2^GFP/+^ adult tendon for CX3CR1 and F4/80 to assess the relationship between these populations during homeostasis. Based on this panel, six different populations or cell states were identified: CCR2+ F4/80+ macrophages (Figure 1E; yellow arrows), CCR2+ CX3CR1+ F4/80+ macrophages (Figure 1E; orange arrows), CCR2-CX3CR1+ F4/80+ macrophages (Figure 1E; pink arrows), CCR2-CX3CR1- F4/80+ macrophages (Figure 1E; red arrows), CCR2-CX3CR1+ F4/80- non macrophages (Figure 1E, white arrows), and a triple negative population. While tissue-resident macrophages only comprise about ∼28% (Figure 1B) of the overall tendon cell environment, these data provide evidence of heterogeneity within the resident macrophage population.

### CCR2 deficiency disrupt macrophage presence during tendon healing

Next, we assessed the presence of CCR2^GFP^+ cells during early and late tendon healing, as well as the impact of CCR2^KO^ on the cellular environment. At 8 days post-repair, CCR2^GFP^ cells comprised 50.57 ± 7.65% of the total cell population in CCR2^GFP/+^ control mice (Figure 2A & B) CCR2^KO^ reduced this population to 10.16 ± 5.66% (Figure 2A & B, p = 0.0020). This significant decrease continued through late healing, with CCR2^GFP^ cells accounting for 45.26 ± 7.65% of all cells in CCR2^GFP/+^ mice, while this population was reduced to 2.53 ± 0.64% in CCR2^KO^ mice at D28 (Figure 2A & B, p=0.0005). Loss of CCR2^GFP^ cells led to a significant reduction in F4/80+ area, relative to controls during early (D8 CCR2^GFP/+^: 33.43 ± 11.82%; D8 CCR2^KO^: 7.38 ± 4.84%, p=0.0072) and late healing (D28 CCR2^GFP/+^: 62.26 ± 17.53%; D28 CCR2^KO^: 19.26 ± 11.37%, p<0.0001). (Figure 2A & C). Consistent with our homeostasis data, CCR2^KO^ healing tendons had reduced numbers of CCR2^GFP^+ F4/80+ macrophages (Figure 2D, yellow arrows), CCR2^GFP^- F4/80+ macrophages (Figure 2D, red arrows) and CCR2^GFP^+ F4/80- cells (Figure 2D, green arrows). Taken together, these data suggest that CCR2^KO^ hinders the presence of CCR2^GFP^+ cells and macrophages during early and late tendon healing.

**Fig 2.**
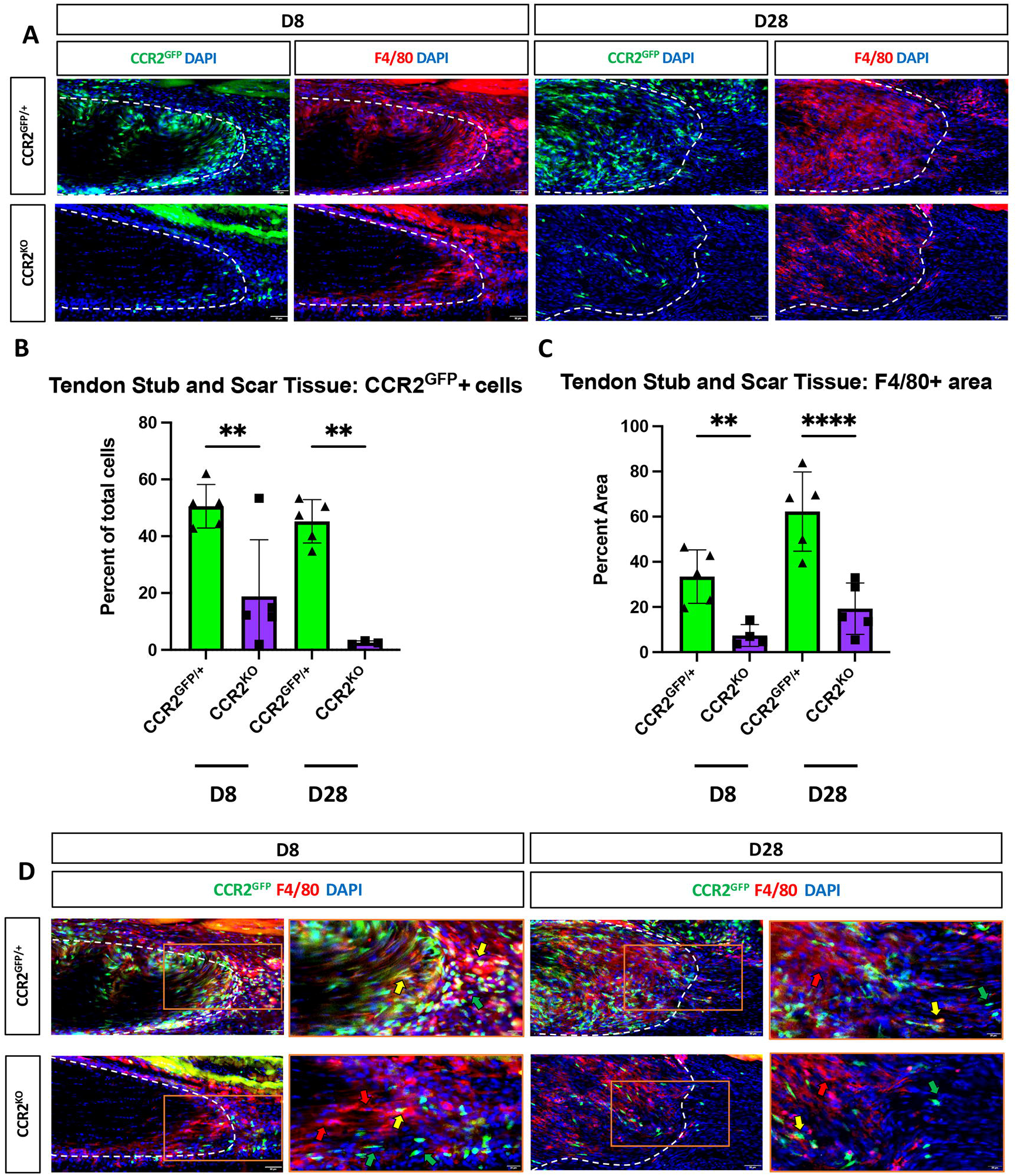
Blunted CCR2 recruitment reduces macrophage presence during early and late healing. (A) To determine the effect of CCR2 deficiency during tendon healing, immunofluorescence of GFP (CCR2^GFP^) and F4/80 at 8- and 28-days post-repair were stained in CCR2^GFP/+^ and CCR2^KO^ healing tendons. Tendon stubs are outlined in dotted white line. (B) Quantification of % of CCR2^GFP^ and (C) % area of F4/80+ macrophages at 8- and 28-days post-repair in CCR2^GFP/+^ and CCR2^KO^ healing tendons. A two-way ANOVA with Sidak’s multiple comparisons test was used to determine statistical significance. *p≤0.05, **p≤0.01, ***p≤0.001. (D) Co-immunofluorescence of GFP (CCR2^GFP^) and F4/80 at 8 and 28 post-repair. Nuclear live cell stain is blue. Scale bars, 50 µM and 20 µM.

### CCR2 deficiency does not affect tendon biomechanics during homeostasis

To determine the functional effects of CCR2^KO^ on tendon homeostasis, contralateral hindlimbs from CCR2^WT^ and CCR2^KO^ mice were harvested at 14-, 21-, 28- and 35-days post-repair for assessment of gliding function and biomechanical analysis. A two-way ANOVA with Sidak’s multiple comparisons test between genotypes and timepoints for the contralateral revealed no significant differences in gliding (gliding resistance and metatarsophalangeal flexion angle) and biomechanical properties (maximum load at failure and stiffness), as such contralateral data from each time-point is presented in aggregate (Figure 3).

**Fig 3.**
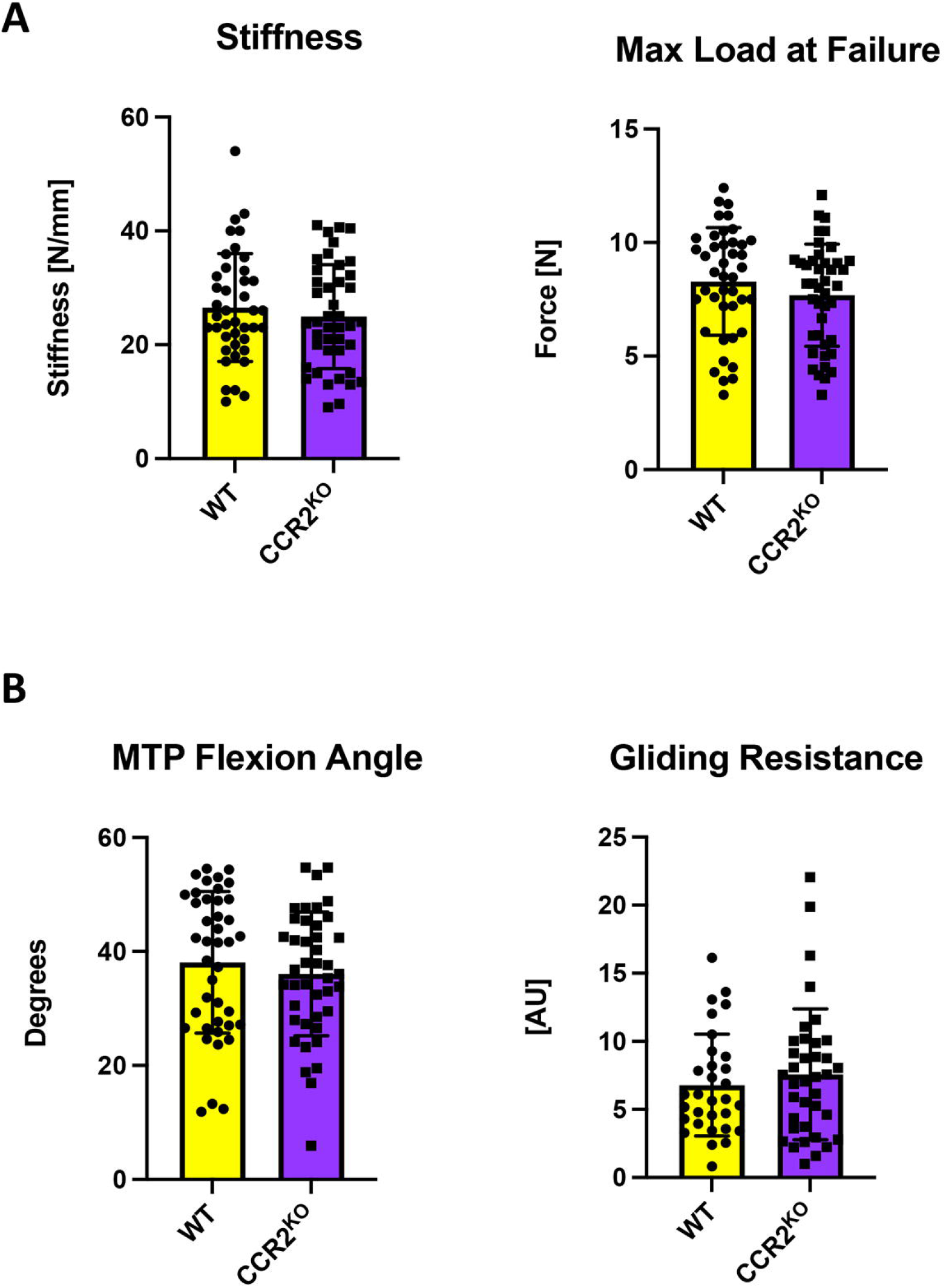
CCR2 deficiency does not affect tendon biomechanics during homeostasis. (A) Measurement of biomechanical properties (stiffness and maximum load at failure) and (B) gliding properties (gliding resistance and metatarsophalangeal joint flexion angle) in uninjured WT and CCR2^KO^ tendons. Student’s *t* test was used to assess statistical significance. A two-way ANOVA with Sidak’s multiple comparisons test between genotypes and timepoints for the contralateral revealed no significant differences in gliding and biomechanical properties, as such contralateral data (n=9-12 per genotype) from each time-point is presented in aggregate. Student’s *t* test was used to assess statistical significance.

### Reduced CCR2+ cell recruitment impedes late tendon healing but does not alter range of motion

To determine the functional effects of CCR2 deficiency during the healing process, CCR2^WT^ and CCR2^KO^ hindlimbs were harvested between 14-35 days post-repair for gliding function and biomechanical analysis. No significant differences in biomechanical properties were observed at D14 and D21 post-repair between CCR2^WT^ and CCR2^KO^ mice. However, at D28, significant decreases in both maximum load at failure (CCR2^WT^: 2.41 ± 0.87 N, CCR2^KO^: 1.29 ± 0.64 N, p = 0.0076) and stiffness (CCR2^WT^: 8.19 ± 4.27 N/mm, CCR2^KO^: 4.39 ± 2.26 N/mm, p= 0.0448) were observed in CCR2^KO^ tendons compared to the control (Figure 4A). By D35 the impairments in maximum load (CCR2^WT^ 3.18 ± 1.04 N, CCR2^KO^: 2.54 ± 0.56 N, p=0.4181) and stiffness (CCR2^WT^: 8.20 ± 2.97 N/mm, CCR2^KO^: 6.91 ± 2.07 N/mm, p=0.9605) in CCR2^KO^ mice were resolved (Figure 4A). Taken together, these data suggest that CCR2^KO^ alters the timing and rate of reacquisition of mechanical properties during tendon healing.

**Fig 4.**
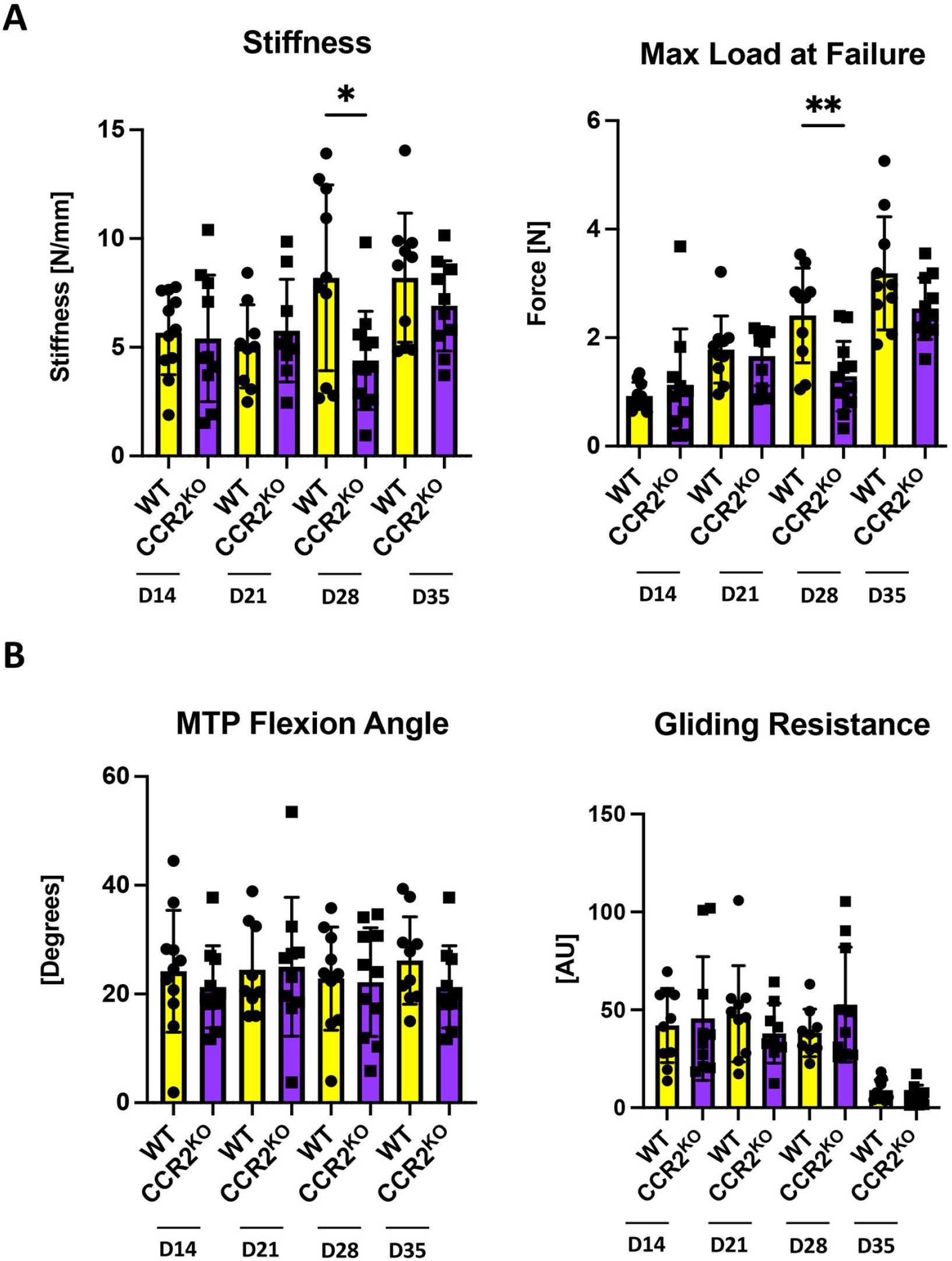
CCR2^KO^ healing tendons impedes late tendon healing. Measurement of (A) biomechanical (stiffness and maximum load at failure) and (B) gliding properties (gliding resistance and metatarsophalangeal join flexion angle) at 14-,21-,28- and 35-days post-repair in WT and CCR2^KO^ tendons. N= 9-12 per genotype/timepoint. Two-way ANOVA with Sidak’s multiple comparisons test was used to assess statistical significance. *p≤0.05, **p≤0.01.

Because macrophages are implicated in modulating scar tissue formation, we also examined the effects of CCR2^KO^ on post-operative tendon range of motion and gliding resistance (Figure 4B). No significant differences in MTP flexion angle or gliding resistance were observed between CCR2^WT^ and CCR2^KO^ tendons at any time-point between 14-35 days.

### CCR2 deficiency reduces myofibroblast content during tendon healing

Because macrophages can influence fibroblast activation and engage in a pro-fibrotic feedback loop with myofibroblasts(11,27), we assessed the effects of CCR2 deficiency on the myofibroblast population during healing. While there was an observable decrease in αSMA staining in CCR2^KO^ tendon repairs at D14 and D28, relative to CCR2^WT^, these differences were not statistically significant (Figure 5B & C; D14: p=0.1151 and D28: p=0.0576). A key step in myofibroblast differentiation is the activation of tissue resident fibroblasts(28). To determine whether CCR2^KO^ may impact this initial step in differentiation, we stained for fibroblast activation protein (FAP) at D14. Indeed, CCR2^KO^ tendons had reduced FAP staining (p=0.0791, Figure 5D & E), relative to CCR2^WT^. Collectively, these data suggest that CCR2^KO^ may impair tenocyte activation and subsequent differentiation to myofibroblasts.

**Fig 5.**
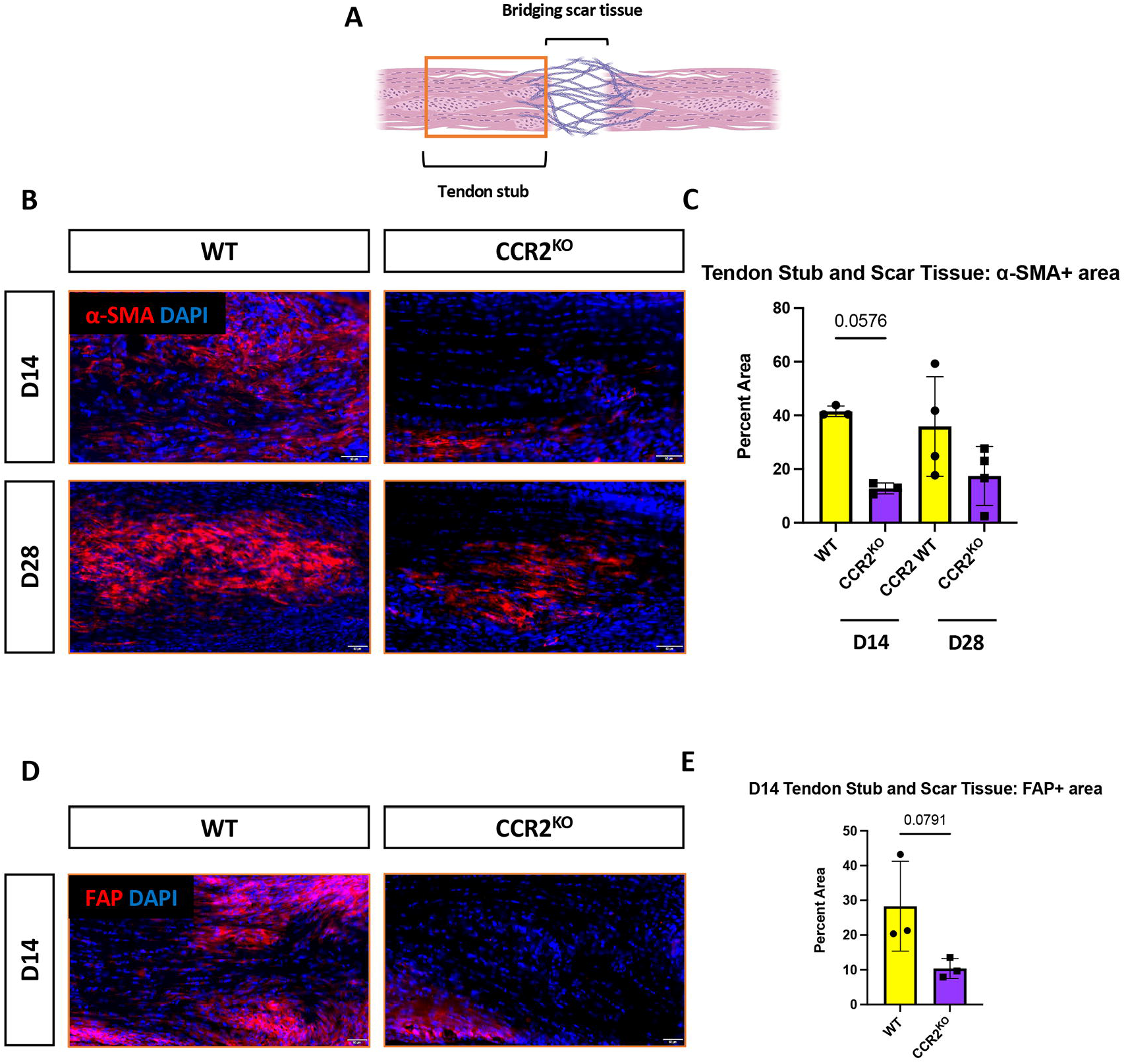
CCR2 deficiency transiently reduces myofibroblast presence. (A) Schematic of fibrotic tendon healing outlining tendon stub and bridging scar tissue. Orange box indicates location of immunofluorescent images. (B) Immunofluorescence staining of α-SMA+ myofibroblasts in WT and CCR2^KO^ tendons stub ends at 14- and 28-days post-repair to define changes in myofibroblasts during CCR2 deficient healing. (C) Quantification of % α-SMA+ area in WT and CCR2^KO^ healing tendon at 14- and 28-days post-repair. Student’s t test was used to assess statistical significance. (D) Immunofluorescence staining of fibroblast activation protein (FAP) in WT and CCR2^KO^ tendon stub ends at 14 days post-repair. (E) Quantification of % area of Fibroblast Activation Protein (FAP) at 14 days post-repair in WT and CCR2^KO^ healing tendons Nuclear live cell stain is blue. Scale bars, 50 µM. N=3-4 per genotype/timepoint.

## DISCUSSION

In the present study, we examined the role of CCR2 in modulating the tendon cell environment during homeostasis and the healing process and tested the hypothesis that loss of CCR2 would be sufficient to obstruct recruitment of extrinsic CCR2+ macrophages. In contrast to this hypothesis, we found that CCR2 also marks tendon-intrinsic cell populations including resident macrophage and T cell populations and that CCR2^KO^ decreases the presence of tendon-resident macrophages during homeostasis. Moreover, CCR2^KO^ decreases the overall macrophage response to tendon injury. Interestingly, we also found that there is CCR2-independent incorporation of macrophages into the tendon during both homeostasis and healing as we observed CCR2^GFP^+ cells present even in CCR2^KO^ mice. In addition to the diminution of the macrophage response, we also observed a reduction in both fibroblast activation and αSMA+ myofibroblast presence during tendon healing. Collectively, these data identify CCR2 as a tendon-resident macrophage and T cell marker, and demonstrate that CCR2^KO^ results in transient impairments during later tendon healing.

Given that CCR2 is predominantly considered a regulator of the release of monocytes from the bone marrow and the recruitment of monocytes to tissues during injury, we were particularly interested in this population during tendon healing. However, we first demonstrated the presence of a CCR2+ tendon resident population, consistent with similar findings in the brain(29), heart(30) and liver(31). While the mechanisms that dictate CCR2+ cell incorporation into homeostatic tissue are unclear, there is some evidence that circulating monocytes replenish tissue resident macrophages during post-natal development via CCR2-mediated mechanisms(32,33). Interestingly, while there was a reduction in CCR2^GFP^ cells in CCR2^KO^ tendons, a complete reduction in this population was not observed, suggesting CCR2-independent mechanisms of cell incorporation as well. The continued presence of CCR2^GFP^ cells in the CCR2^KO^ tendons is consistent with other types of injury using this model(31,34,35), in which CCR2^KO^ results in ∼80% reduction in CCR2 cells, likely because the remaining cells are being recruited in a CCR2-independent manner(36).

The relationship between CCR2+ cells and other known macrophage populations in the tendon during both homeostasis and healing is important to further unravel the complex cell environment that regulates maintenance of tendon health and physiological tendon healing. For example, CX3CR1+ cells have been proposed to be a macrophage-like tendon cell population, or ‘tenophage’, that regulates proliferative and fibrotic responses and has vasculoprotective functions(8). Here we show that while there is some overlap between CCR2+ and CX3CR1+ cells, there are distinctly separate CCR2+ and CX3CR1+ populations, further demonstrating the molecular complexity of the resident tendon cell environment. Finally, the finding of both CCR2+ and CCR2-tissue resident macrophages will be an important area of future work as cardiac resident CCR2+ macrophages express higher levels of inflammatory cytokines and act as critical drivers of monocyte and neutrophil recruitment, while CCR2-macrophages express several growth factors and enhances macrophage proliferation(30). Additionally, recent single cell sequencing data has defined three distinct murine resident macrophage subpopulations across several organs, emphasizing resident macrophage heterogeneity(37).

Consistent with our single cell RNA sequencing data, CCR2 expression has also been reported in regulatory T cells(38) and Th17 cells(39). Regulatory T cells (Tregs) are crucial to the maintenance of tissue homeostasis by preventing autoimmunity and controlling excessive inflammation(40). CCR2^-/-^ T cells promoted accumulation of Tregs and decreased levels of Th17 cells(41), supporting an important role for CCR2 in modulating the T cell response. Recent work has only begun to appreciate the role of T cells in tendon(42,43). Moreover, while a lack of B and T cells did not disrupt tendon development(5), adoptive transfer of neonatal Tregs into adult hosts improved Achilles tendon functional recovery and polarized macrophages into an anti-inflammatory profile(44). Collectively, these data support an important role for T cells in the healing process, and merit future work to better define the role of these cells in the healing process.

Given the potential for macrophages and myofibroblasts to engage in a pro-fibrotic feedback loop in many tissues(11,27), we were particularly interested in the impact of CCR2^KO^ on the myofibroblast environment. In response to macrophage-derived cytokines, fibroblasts undergo an activation process and subsequent myofibroblast differentiation. Moreover, myofibroblast activity can sustain macrophage function, creating an effective pro-fibrotic loop. Prior work has shown that CCR2^KO^ reduces myofibroblast differentiation in injured kidneys(19). Consistent with these data and given the role of macrophages in wound debridement and myofibroblasts in ECM remodeling, reductions in these cell populations and decreased fibroblast activation are a potential culprit of the functional deficit we see during late tendon healing (D28). However, it is not entirely clear why CCR2^KO^ results in only transient mechanical deficits, such that by D35 CCR2^KO^ and CCR2^GFP/+^ tendons are not significantly different. One potential mechanism may relate to different myofibroblasts dynamics during healing. While CCR2^KO^ tendons healed with reduced myofibroblast content at D14, by D28 these differences have resolved so it is possible that restoration of the myofibroblast environment by D28 is sufficient to drive functional restoration by D35. In addition, it remains unknown whether it is the reduction in tissue resident or circulating CCR2+ cells that leads to decreased fibroblast activation and myofibroblast content, and subsequent delays in functional recovery in CCR2^KO^ tendon repairs.

While this study clearly demonstrates that CCR2^KO^ alters the composition of the basal tendon cell environment and the tendon healing process, there are a few limitations that must be considered. This CCR2^KO^ model results in global and sustained deletion of CCR2, and thus global and sustained alterations in macrophage recruitment. Given that macrophages are a highly plastic cell type that can change in response to their microenvironment, depletion of macrophages likely results in altered responsiveness of other cell populations. Future work using temporally-controlled CCR2 inhibition will be important to define the function of these cell during different phases of the healing process. Furthermore, lack of specific markers for tendon-resident and extrinsic macrophages complicates our ability to delineate the role of these cells during healing.

Taken together, our results suggest that CCR2 identifies tendon resident macrophage and T cell populations and that CCR2^KO^ alters the composition of the basal tendon-resident cell environment. In addition, hindered recruitment of CCR2+ cells decrease the macrophage response to tendon injury as hypothesized, with decreased macrophage response during tendon healing leading to decreased myofibroblast content and a subsequent transient slowing of the reacquisition of mechanical properties after tendon injury. As such, these data suggest that a combination of altering the tissue resident and extrinsic macrophage response can drive impaired tendon healing; however, parsing out the relative contributions of these populations, as well as their time-dependent functions will be critical to informing therapeutic target selection moving forward.

## Non-standard abbreviations

αSMA: alpha smooth muscle actin
CCR2: Chemokine Receptor 2
CCR2^-/-^: CCR2 knockout mice
GFP: Green Fluorescent Protein
FAP: Fibroblast Activation Protein
FDL: Flexor Digitorum longus

## Data Availability Statement

The data that support the findings are available in the methods of this article.

## Acknowledgements

We would like to thank the Histology, Biochemistry and Molecular Imaging (HBMI), the Biomechanics, Biomaterials and Multimodal Tissue Imaging (BBMTI), and the Multiphoton Imaging Cores for technical assistance. This work was supported in part by NIH/ NIAMS R01 AR077527 and R01AR073169 (to AEL), NIH/NIAMS T32 AR076950 and NIH/NIAMS K99 AR080757 (to AECN). The HBMI and BBMTI Cores are supported by NIH/ NIAMS P30AR069655.

## Author contributions

Study conception and design: SNM, AEL; Acquisition of data: SNM, AECN, EG; Analysis and interpretation of data: SNM, AECN, AEL; Drafting of manuscript: SNM, AEL; Revision and approval of manuscript: SNM, AECN, EG, AEL.

